# Peptidyl tRNA hydrolase is required for robust prolyl-tRNA turnover in *Mycobacterium tuberculosis*

**DOI:** 10.1101/2022.11.15.516451

**Authors:** Francesca G. Tomasi, Jessica T. P. Schweber, Satoshi Kimura, Junhao Zhu, Laura A. T. Cleghorn, Susan H. Davis, Simon R. Green, Matthew K. Waldor, Eric J. Rubin

## Abstract

Enzymes involved in rescuing stalled ribosomes and recycling translation machinery are ubiquitous in bacteria and required for growth. Peptidyl tRNA drop-off is a type of abortive translation that results in the release of a truncated peptide that is still bound to tRNA (peptidyl tRNA) into the cytoplasm. Peptidyl tRNA hydrolase (Pth) recycles the released tRNA by cleaving off the unfinished peptide and is essential in most bacterial species. We developed a sequencing-based strategy called Cu-tRNAseq to study the physiological role of Pth in *Mycobacterium tuberculosis* (*Mtb*). While most peptidyl tRNA species accumulated in a strain with impaired Pth expression, peptidyl prolyl-tRNA was particularly enriched, suggesting that Pth is required for robust peptidyl prolyl-tRNA turnover. Reducing Pth levels increased *Mtb*’s susceptibility to tRNA synthetase inhibitors that are in development to treat tuberculosis (TB) and rendered this pathogen highly susceptible to macrolides, drugs that are ordinarily ineffective against *Mtb*. Collectively, our findings reveal the potency of Cu-tRNAseq for profiling peptidyl tRNAs and suggest that targeting Pth would open new therapeutic approaches for TB.

## Introduction

Enzymes involved in ribosome rescue are ubiquitous in bacteria [1]. mRNA truncation, frame-shifting, and readthrough errors can result in unproductive ribosomes that stall on transcripts because the stop codon at the end of mRNA is missing or no longer in-frame [2]. Certain amino acid combinations or codon patterns can also interfere with ribosome efficiency and lead to stalling [3]. Even in the absence of cellular stressors, hiccups in protein synthesis occur with sufficient frequency that the accumulation of stalled ribosomes would impair cell survival [4].

To rescue stalled ribosomes, bacteria use a diverse set of pathways [5] which all result in stalled ribosomes being returned to the free pool of actively translating ribosomes. One mechanism of freeing the ribosome is peptidyl tRNA drop-off, or simply ‘drop-off’, where peptidyl tRNA dissociates from the ribosome either spontaneously or with the help of ribosome recycling factor (RRF), elongation factor G (EF-G), and release factor 3 (RF3) [6–8]. A key distinction in drop-off is the release of peptidyl tRNA into the cytoplasm; by contrast, rescue systems like trans-translation and ArfA/B hydrolyze peptidyl tRNA that is still bound to the ribosome using different enzymes [1].

The central enzyme in peptidyl tRNA drop-off is peptidyl tRNA hydrolase (Pth), an esterase that recycles tRNA by cleaving the released peptidyl tRNA [7]. While the structure of Pth has been solved in multiple organisms and its catalytic mechanism of action is well understood, little is known about the functional roles of peptidyl tRNA drop-off in bacteria in vivo [9, 10]. tRNA turnover is complex and tightly regulated in cells [11], and Pth’s role in tRNA dynamics is not well-understood. Pth does not have differential activity on different tRNA species *in vitro* [12–14], but it is unclear whether this enzyme preferentially cleaves specific tRNA iso-acceptors *in vivo*.

Here, we examine the physiological role of Pth in *Mycobacterium tuberculosis* (*Mtb*), a pathogen that was responsible for over 10.5 million active cases of tuberculosis (TB) and at least 1.5 million deaths in 2020 [15]. *Mtb* has been shown to be highly susceptible to the inhibition of transtranslation during normal growth, suggesting this pathogen has a critical need for ribosome rescue machinery [16]. Work on trans-translation in *Mtb* has opened doors for drug discovery efforts aimed at targeting ribosome rescue in *Mtb* [17]. However, peptidyl tRNA drop-off and the effects of Pth depletion in *Mtb* have not yet been investigated. Here we have developed Cu-tRNA sequencing to profile *Mtb* peptidyl tRNAs and to study Pth in *Mtb*. We show that while peptidyl tRNA accumulates across most charged tRNA species in a Pth hypomorph, peptidyl prolyl-tRNA overtakes proline tRNA pools in a Pth knockdown, suggesting that Pth is required for robust prolyl-tRNA turnover. Reducing Pth levels increased *Mtb’s* susceptibility to tRNA synthetase inhibitors and macrolides, antibiotics that are not currently used to treat TB. Our work underscores the importance of Pth in tRNA turnover and opens new avenues for anti-TB strategies that exploit synergy between translation errors and tRNA turnover.

## Results

### *pth* is required for normal growth of *Mtb*

Transposon sequencing in *Mtb* suggests that, like in *E. coli*, the gene encoding Pth (Rv1014c) is required for survival [18, 19]. We constructed two types of *Mtb* Pth mutants using complementary genetic techniques: proteolytic degradation and CRISPR interference (CRISPRi). While the first approach depletes intracellular Pth levels by tagging proteins for degradation using a titratable system [20], the other achieves knockdown through transcriptional repression of varying strengths based on the protospacer-adjacent motif (PAM) used to target catalytically dead Cas9 (dCas9) to *pth* [21]. Both types of strains had growth defects that were proportional to the level of depletion (Figure 1), confirming that *pth* is required for normal growth of *Mtb*. Due to the technical ease of validating *pth* knockdown levels and our use of the tightly-regulated transcriptional repression strain in published work [22], we used the CRISPRi Pth depletion strain for subsequent experiments in this work.

**Figure 1.**
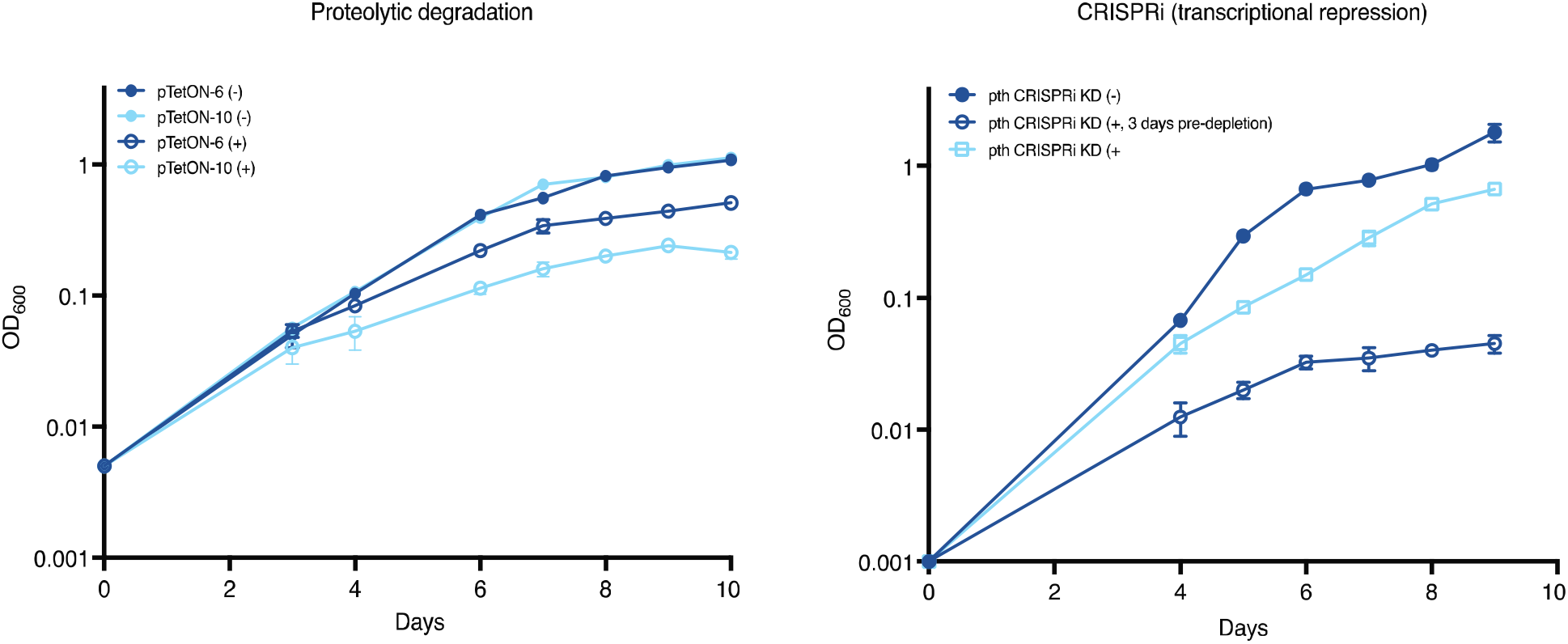
Complementary techniques show Pth is required for normal growth in *Mtb*. Growth curves are shown for different Pth knockdown constructs: proteolytic degradation (left) and transcriptional repression (right). In both panels, (-) = uninduced and (+) = induced for Pth knockdown. Growth was measured spectrophotometrically for each strain with OD_600_ measurements over the course of 10 days in the presence or absence of inducer. Prior to taking measurements, strains were diluted to the starting OD_600_ indicated at day 0. Numbers next to the plasmid name indicate promoter strength for SspB expression (6 = lower, 10 = higher, corresponding to the expected relative level of protein depletion). “Pre-depletion” refers to growing cells in the presence of aTC for 3-4 days prior to diluting strains to the indicated starting OD_600_. Predepletion enables a higher level of *pth* knockdown to be achieved before conducting experiments.

We previously characterized a second function for Pth in *Mtb*. In addition to its peptidyl tRNA hydrolase activity, Pth is a detoxifying enzyme for a tRNA-acetylating toxin TacT that is part of a newly-identified toxin-antitoxin system [22]. We found that this function does not contribute to *pth* essentiality in *Mtb* since TacT was not active during normal laboratory growth conditions and a *tacAt* knockout strain was equally susceptible to *pth* depletion by CRISPRi [22]. Thus, we focused our efforts on studying the effects of Pth depletion in *Mtb* on peptidyl tRNA pools and tRNA turnover following translation errors.

### Cu-tRNAseq measures peptidyl tRNA levels

Studies of *E. coli* have shown that Pth depletion leads to the accumulation of peptidyl tRNA in cells [13]. The identification of peptidyl tRNAs has typically been accomplished using Northern blotting [23], making it technically difficult to compare pools across all tRNA species and their respective iso-acceptors. More recent work on other substrates for Pth such as N-acetylated aminoacyl tRNAs has surveyed tRNA species using mass spectrometry, which substantially increases sensitivity but still poses scalability challenges [22, 24, 25]. We developed a tRNA sequencing-based strategy, dubbed Cu-tRNA-seq, to study the effects of *pth* depletion on peptidyl-tRNA profiles in a rapid, quantitative, and high-throughput manner (Figure 2).

**Figure 2.**
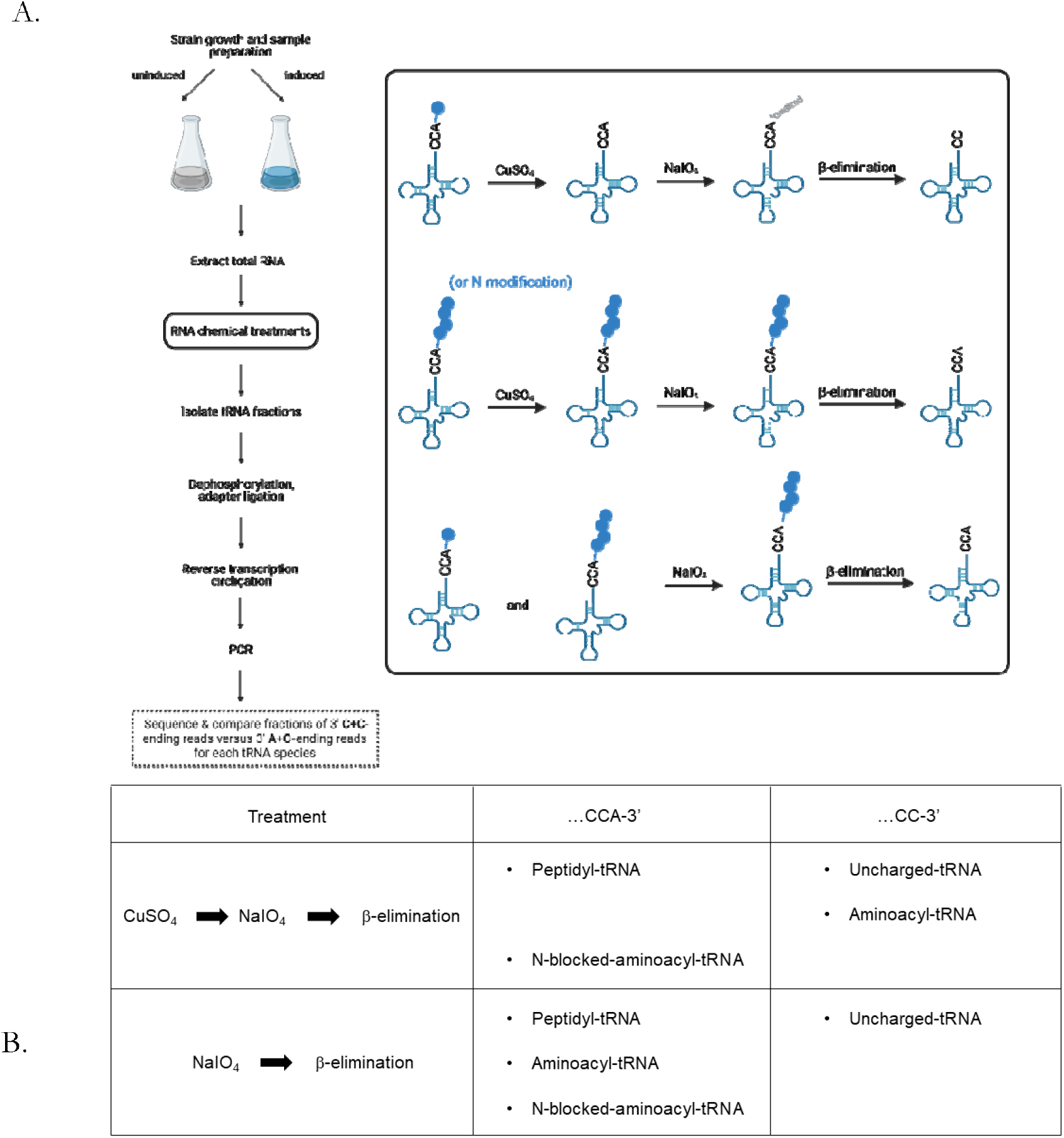
Application of tRNA sequencing to measure pools of peptidyl and N-acetylated tRNA. Overview of the copper sulfate-based tRNA sequencing (Cu-tRNAseq) protocol. Strains are grown and harvested at a normalized cell density. Total RNA is subjected to the chemical treatments (boxed) to distinguish between aminoacyl or uncharged tRNA and peptidyl or N-blocked (e.g., N-acetylated) aminoacyl tRNA (top and middle, respectively). A control treatment was performed for each replicate without CuSO_4_ to measure total charged tRNA levels (bottom). tRNA fractions are extracted from 1-2 μg of total RNA for sequencing library preparation as described in Methods [26]. (B) The ratio of 3’ A+C-ending reads compared to total 3’C+C and 3’A+C ending reads for each tRNA is computed as described [27] to estimate the fraction of peptidyl (or N-acetylated) tRNA in each tRNA species. A higher fraction of A+C-ending reads following Cu-tRNAseq suggests a higher proportion of peptidyl or N-acetylated aminoacyl tRNA. Figure created with BioRender.com.

tRNA sequencing has been used to survey tRNA levels and modifications in both bacteria and eukaryotes and recently an application was developed to measure aminoacyl-tRNA fractions [27]. This method incorporates chemical treatments prior to sequencing that distinguish between charged and uncharged tRNA. All tRNAs share a 3’ CCA tail that serves as the aminoacylation site [28]. The first chemical treatment – oxidation by sodium periodate – has no effect on the 3’ end of a charged tRNA, because the bound amino acid protects molecules from oxidation. However, an uncharged tRNA is oxidized at the 3’A. A subsequent β-elimination reaction at high pH then removes an unprotected 3’ A residue at the end of uncharged tRNA, or simply deacylates charged tRNA, leaving the 3’A intact on the molecule. Sequencing of tRNA libraries and comparing CC-ending reads to CCA-ending reads for each tRNA isoacceptor provides a global view of tRNA molecules that are protected by single amino acids or peptides at their 3’ ends [27].

We adapted this technique to monitor peptidyl tRNAs by adding a chemical treatment with copper sulfate (CuSO_4_) prior to periodate oxidation (Figure 2). Incubation with CuSO_4_ deacylates tRNAs charged with a single amino acid, but this treatment leaves N-blocked aminoacyl tRNA (such as peptidyl tRNA, N-acetylated aminoacyl tRNA, or formyl-methionine tRNA) untouched (Figure 2A) [23, 29]. Thus, in sequencing data, peptidyl tRNA and other N-blocked aminoacyl-tRNAs will have intact CCA 3’ ends, whereas aminoacyl-tRNAs and deacylated tRNAs will have CC 3’ ends (Figure 2B).

As proof of principle for the Cu-tRNAseq approach, we used our previously described *M. smegmatis* (*Msmeg*) strain that overexpresses *Mtb* TacT, a toxin that adds an acetyl group to the amino acid on charged tRNA [22]. *Msmeg* lacks *tacT*, and since *Mtb* TacT acetylates glycyl-tRNA, we hypothesized that the protected glycyl-tRNA fraction would increase with TacT overexpression. Indeed, we observed an increased percentage of protected glycyl-tRNA (Figure 3), suggesting that Cu-tRNAseq can be used to measure N-blocked aminoacyl tRNA. Additional support for this conclusion was the observation that there were relatively high levels of protected initiator tRNA (fMet), which contains a formylated methionine. Histidyl-tRNA was also resistant to CuSO_4_ deacylation, suggesting that histidine’s unique imidazole ring affects the reaction’s efficiency. Together, these observations indicate that Cu-tRNAseq provides a method to distinguish N-blocked aminoacyl tRNAs from aminoacyl tRNAs and therefore should be useful for measuring peptidyl tRNA levels.

**Figure 3.**
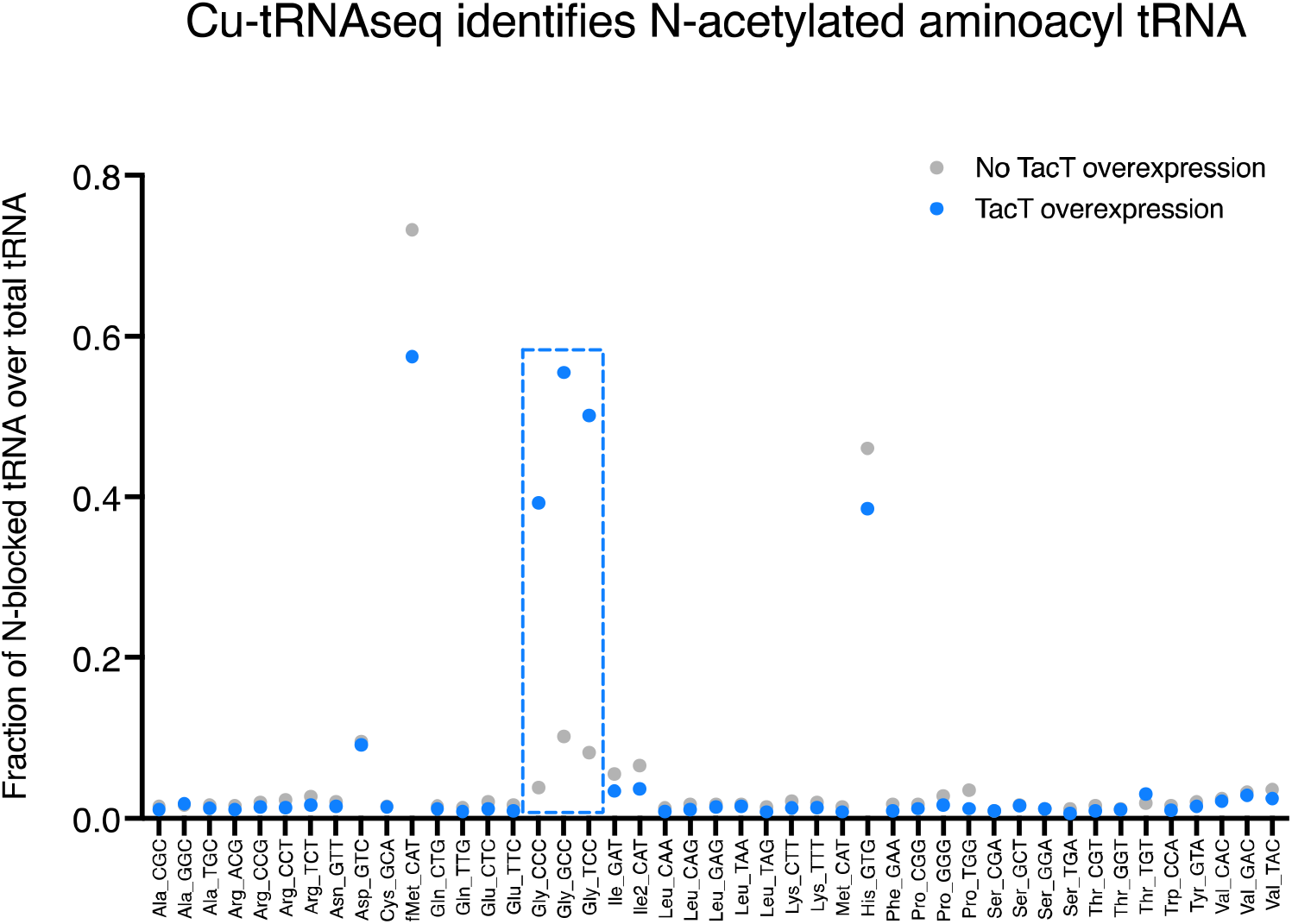
CuSO_4_-tRNA sequencing (Cu-tRNAseq) distinguishes N-acetylated aminoacyl tRNA from aminoacyl or uncharged tRNA. Cu-tRNAseq was carried out on a previously described *M. smegmatis* (*Msmeg*) strain with inducible *Mtb* TacT overexpression [22]. The fraction of peptidyl tRNA is plotted as a ratio of the number of 3’CCA-ending reads to the sum of 3’CCA- and 3’CC-ending reads for each tRNA species. Cu-tRNAseq detected N-acetylated glycyl-tRNA in the induced (TacT overexpression) condition (blue dashed box).

### Pth depletion in *Mtb* reduces pools of usable tRNA

We applied Cu-tRNAseq to *Mtb* samples to compare tRNA profiles between strains that were induced for *pth* depletion by CRISPRi and uninduced strains with wild-type Pth levels. The three chemical pre-treatments, CuSO_4_, NaIO_4_, and β-elimination, as well as Pth depletion did not appreciably skew tRNA read counts (Table S1, Figure S1).

Without pth depletion, in most tRNA species, a small fraction was protected after copper sulfate treatment, consistent with the idea that generally only a tiny fraction of tRNA exists as peptidyl-tRNA. However, some tRNA species, such as tRNA-Asp, -fMet, -His, and -Ile showed a higher proportion of protected species, suggesting that these peptidyl-tRNAs accumulated or that these aminoacyl-tRNAs are more tolerant to CuSO_4_ treatment (as is the case with fMet). With Pth knockdown, the proportion of tRNAs protected from copper sulfate treatment generally increased (Figure 4A). The accumulation of protected proline tRNAs was particularly dramatic, suggesting that there is marked accumulation of peptidyl prolyl-tRNAs in the absence of wild type Pth levels. Acid-PAGE northern blotting was used to corroborate that peptidyl prolyl-tRNA accumulates in these conditions (Figure S2). These results indicate that peptidyl prolyl-tRNAs specifically accumulate when Pth is depleted, suggesting that Pth promotes robust turnover of peptidyl prolyl-tRNAs in wild-type Mtb cells.

**Figure 4.**
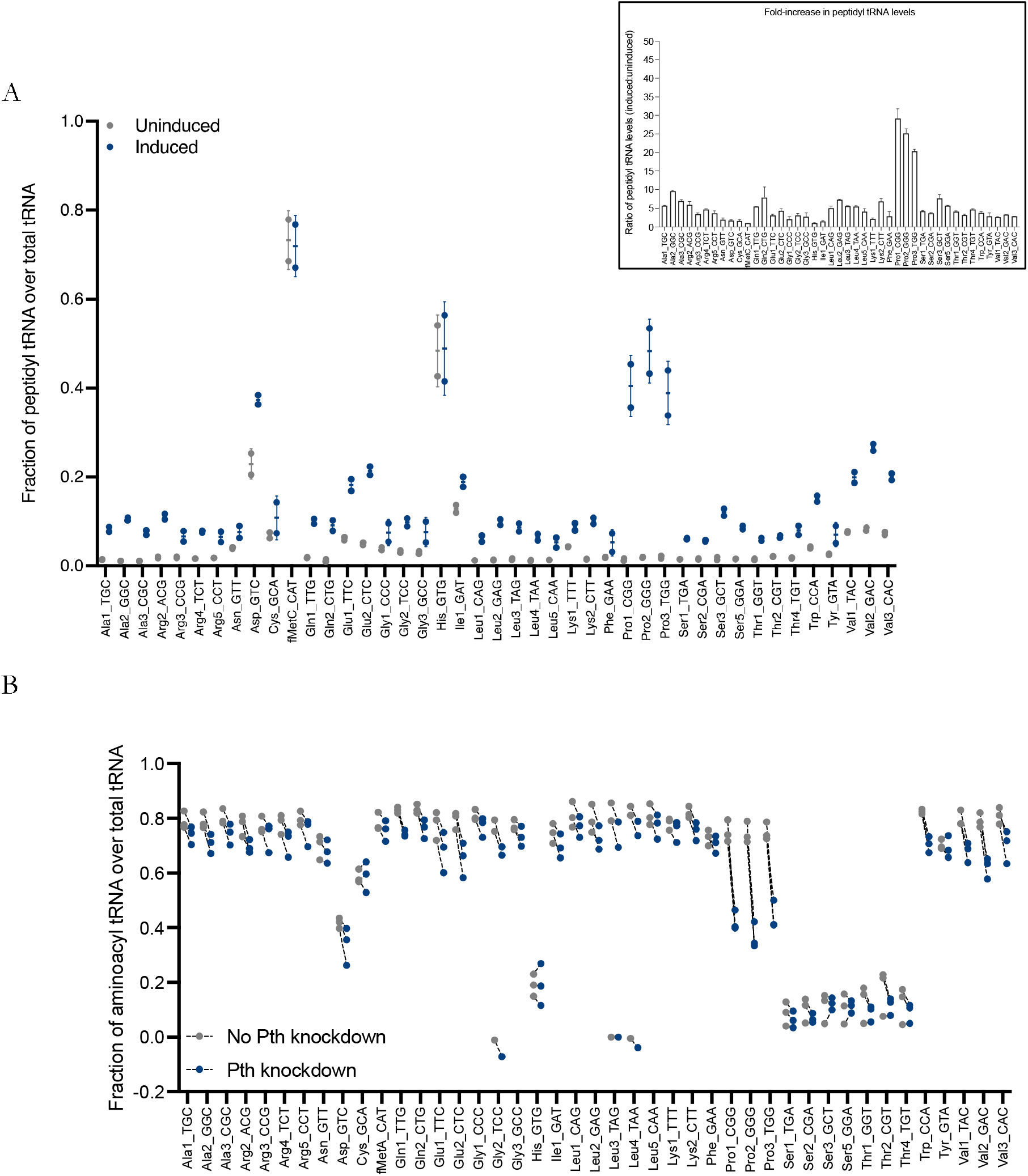
Depleting Pth leads to accumulation of peptidyl tRNAs in *Mtb* and a reduction in usable aminoacyl tRNA pools. Cu-tRNAseq was carried out in a *Mtb* CRISPRi Pth knockdown. (A) Strains were grown to an OD_600_ of 0.3-0.4 in the presence or absence of inducer (50 ng/mL aTC) and total RNA was harvested. The fraction of peptidyl tRNA is calculated as the ratio of the number of 3’CCA-ending reads to the sum of 3’CCA- and 3’CC-ending reads for each tRNA species. The bar plot (top right) displays the ratio of peptidyl tRNA fractions between induced and uninduced strains. (B) RNA samples from (A) were treated with periodate and ß-elimination only, without CuSO_4_ (Dataset S2) to measure the fraction of the tRNAs protected by either a single amino acid or a peptide. The aminoacyl tRNA fraction for each species is calculated by subtracting the protected fraction with CuSO_4_ treatment from the total protected fraction. Dashed lines connect protected tRNA fractions in uninduced (no Pth knockdown) with induced (Pth knockdown) samples to show an overall decrease in usable charged tRNA pools. Serine and threonine tRNAs are typically acylated at lower levels than other tRNAs in bacteria [30].

We hypothesized that the accumulation of peptidyl-tRNAs would be accompanied by a decrease in aminoacyl-tRNA levels. To profile aminoacyl tRNA levels, we conducted tRNA-seq with periodate treatment and β-elimination only and measured the ratio of the protected fractions, which together represent the total of aminoacyl-tRNA and peptidyl-tRNA fractions. The aminoacyl-tRNA proportion was calculated by subtracting the proportion of peptidyl-tRNA fractions measured by Cu-tRNAseq (Figures 4A and B). Prolyl-tRNA availability was drastically decreased in the Pth depletion condition due to the accumulation of peptidyl prolyl-tRNA, indicating that Pth is required for maintaining available prolyl-tRNAs in normal conditions (Figure 4B).

### Pth depletion increases *Mtb* sensitivity to candidate aminoacyl tRNA synthetase inhibitors

Since the tRNA sequencing findings revealed that Pth promotes tRNA turnover in *Mtb*, we hypothesized that further targeting a single tRNA species would have a compound effect on *Mtb* survival in the absence of Pth; i.e., we explored whether limiting tRNA availability by some other mechanism would synergize with Pth depletion. Drugs that inhibit aminoacyl tRNA synthetases have seen clinical success against malaria (halofuginone) and certain skin infections (mupirocin) and efforts are currently underway to develop similar inhibitors for TB therapeutics. We tested a candidate lysine tRNA synthetase inhibitor and found some synergy with Pth knockdowns compared to uninduced controls (Figure 5). These observations suggest that orthogonally targeting *Mtb*’s ability to maintain normal aminoacyl tRNA levels could represent a powerful anti-TB strategy and hypothesize that synergy would be particularly potent with a candidate proline tRNA synthetase inhibitor, since lysine tRNA levels were less drastically impacted by a Pth knockdown than proline tRNA.

**Figure 5.**
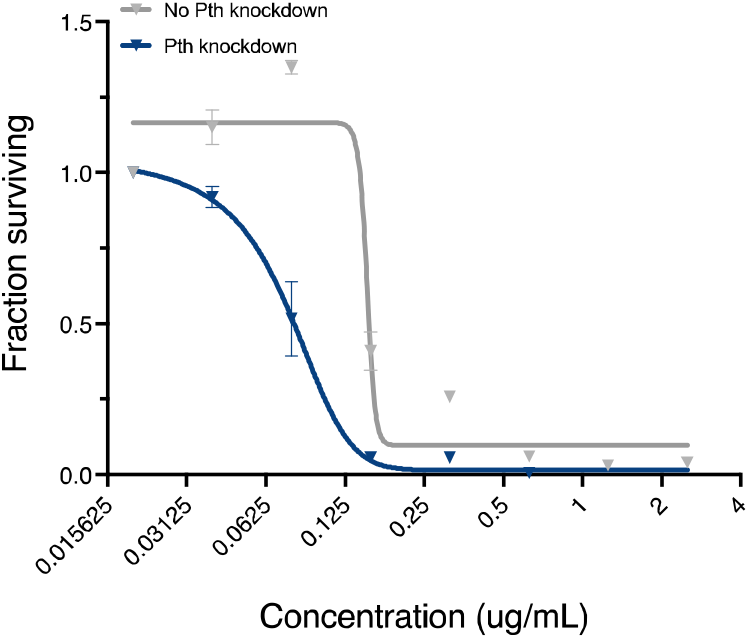
Pth depletion increases *Mtb* sensitivity to a candidate lysine tRNA synthetase inhibitor. *Mtb* growth in the presence a candidate lysine tRNA synthetase inhibitors [31] is plotted across drug concentrations as determined by fluorescence in an Alamar blue assay. Pth CRISPRi strains were grown to mid-log and diluted to an OD_600_ of 0.001 and plated in serial dilutions of antibiotic in 96-well plates. The fraction of *Mtb* cells surviving is plotted by normalizing fluorescence values to control wells with no drug, and a least squares fit of dose response data is plotted. Results are representative of two biological replicates.

### Pth depletion sensitizes *Mtb* to macrolide antibiotics

In *E. coli*, a temperature-sensitive Pth mutant is hypersusceptible to macrolides, a class of antibiotics that target the large 50S ribosomal subunit [32]. Macrolides are thought to increase rates of peptidyl tRNA drop-off by obstructing the nascent peptide exit tunnel and leading to early ribosome dissociation from transcripts [33, 34]. Macrolides are not currently used to treat tuberculosis but are generally well-tolerated and are used to treat non-tuberculous mycobacterial infections [35–37]. To explore whether inhibition of Pth would sensitize *Mtb* susceptible to macrolides, we tested our collection of Pth knockdown strains against a panel of macrolides. Interestingly, all levels of Pth depletion tested rendered *Mtb* hypersusceptible to macrolides but not to other ribosome-targeting antibiotics that have different mechanisms of action (Figure 6A, Table S2). Pth hypomorphs were also more susceptible to the lincosamide clindamycin and the aminonucleoside puromycin, both of which also trigger peptidyl tRNA drop-off (Figure S3) [32, 38]. These results suggest that inhibitors that partially block wild-type Pth activity would render *Mtb* susceptible to treatment with macrolides or other drugs that also increase peptidyl tRNA drop-off.

**Figure 6.**
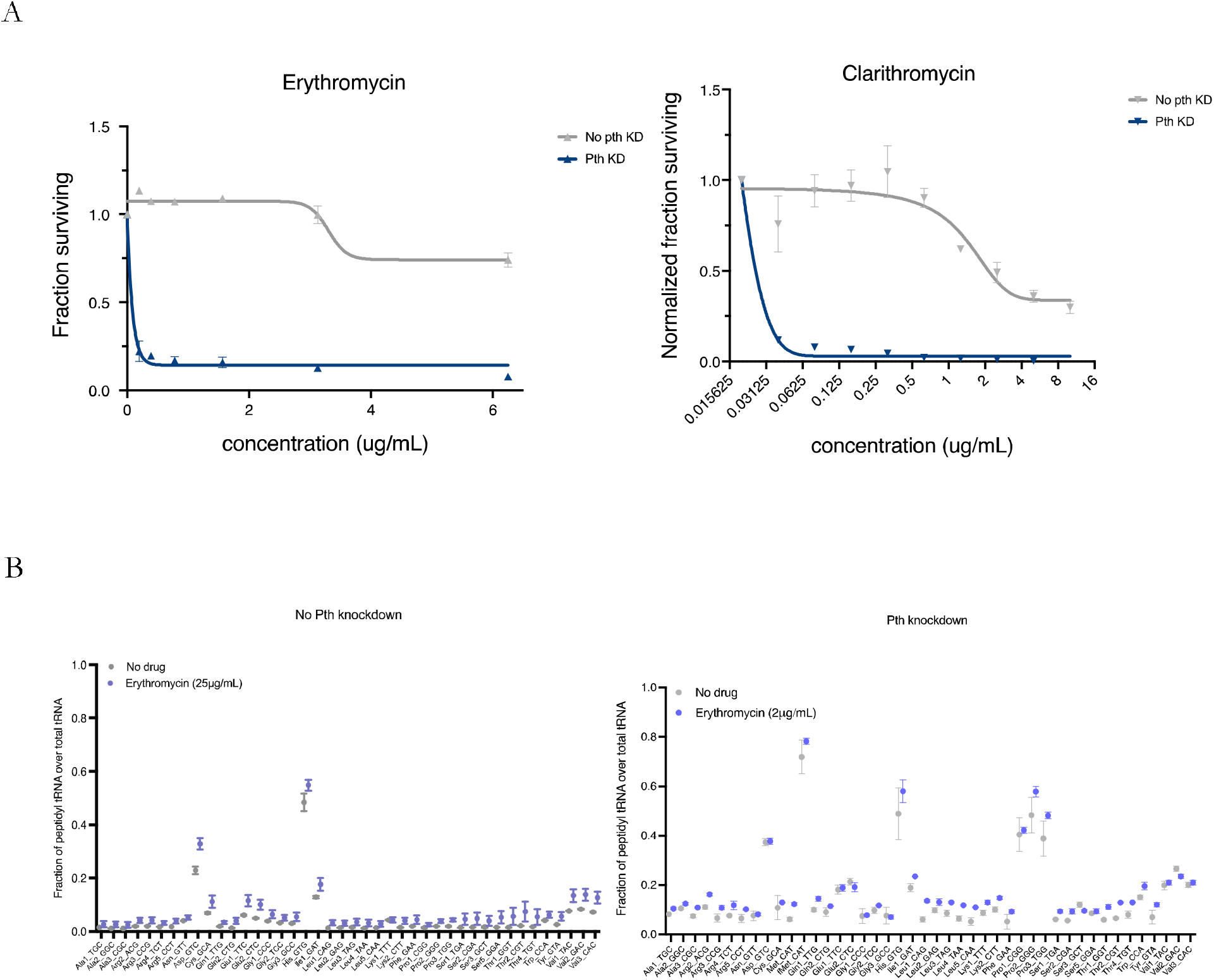
Pth depletion sensitizes *Mtb* to macrolides. (A) *Mtb* growth in the presence of two macrolides – erythromycin and clarithromycin – is plotted across drug concentrations as determined by fluorescence in an Alamar blue assay, and a least squares fit of dose response data is plotted [39]. Pth CRISPRi strains were grown to mid-log and diluted to an OD_600_ of 0.001 and plated in serial dilutions of antibiotic in 96-well plates. The fraction of *Mtb* cells surviving is plotted by normalizing fluorescence values to control wells with no drug. Results are representative of three biological replicates. (B) Cu-tRNAseq was performed on Pth CRISPRi strains in the presence and absence of erythromycin. Strains were grown in the presence or absence of inducer as described in Figure 3. Uninduced strains were then treated overnight with 25 μg/mL of erythromycin, and Pth knockdown-induced strains were treated with 2 μg/mL for the same duration. Results are from three biological replicates.

Indeed, tRNA sequencing shows a slight accumulation in wild type *Mtb* treated with an inhibitory concentration of erythromycin, and a bigger increase in peptidyl tRNA pools across tRNAs in a Pth knockdown treated with erythromycin (Figure 6B). The effect in wild type cells is likely negligible owing to the presence of Pth, but the larger fractions of peptidyl tRNA in the Pth hypomorph suggests that cells are overloaded with Pth substrates upon erythromycin treatment. This effect was not observed on tRNA pools from *Mtb* treated with chloramphenicol (Figure S4). These findings confirm that macrolides increase peptidyl tRNA drop-off, and that even slightly perturbing tRNA pools in a Pth knockdown leads to significant phenotypic consequences.

## Discussion

In this work, we developed a new application of tRNA sequencing to monitor peptidyl tRNA levels. Cu-tRNAseq builds on existing techniques by incorporating copper sulfate treatment to chemically distinguish peptidyl tRNA from aminoacyl or uncharged tRNA, facilitating the study of the composition of tRNA pools in a quantitative and high-throughput manner. Using Cu-tRNAseq, we found that depleting Pth led to a reduction in most aminoacyl tRNAs but that proline tRNA pools were the most substantially impacted. Our findings suggest that Pth is particularly important for recycling peptidyl tRNA that has dropped off during the incorporation of proline to a growing peptide in *Mtb*. Mycobacteria are a highly GC-rich genus, and pathogenic mycobacteria are enriched for so-called PE/PPE proteins, outer membrane proteins characterized by their proline-rich N-terminal sequences that make up nearly 10% of the *Mtb* genome [40, 41]. It is tempting then to speculate that *Mtb*’s proline richness contributes to a critical need for Pth to promote robust proline tRNA turnover and keep up with cells’ demand for prolyl-tRNA.

Peptidyl tRNA drop-off early on in translation is thought to be mediated by the ribosome release and recycling factors RF3 and RRF [6]. While the peptidyl tRNA substrates from this mechanism of drop-off have not specifically been characterized prior to this work, recent studies have characterized another phenomenon known as intrinsic ribosome destabilization (IRD), in which a longer nascent chain can also trigger premature translation termination [42]. While the mechanism of IRD also requires further study, peptidyl tRNA released into the cytoplasm during abortive IRD can be a substrate for Pth [42]. Interestingly, studies of these truncated peptides found a relationship between the presence of alternating proline and acidic amino acids and the likelihood that a ribosome will be destabilized [42]. These findings corroborate our findings that proline amino acids predispose ribosomes to abortive translation.

Even though proline tRNA turnover was the most impacted by Pth depletion, we found that *Mtb* Pth hypomorphs were more susceptible to lysine tRNA synthetase inhibitors than uninduced controls. This suggests that the smaller reductions we observed in aminoacyl tRNA availability in other tRNAs (where there were lower rates of peptidyl tRNA drop-off) may also be biologically significant, *pth* is essential in many other bacterial species including *E. coli* and *B. subtilis*, neither of which is enriched for polyproline containing proteins, suggesting that Pth controls tRNA pools more broadly. In fact, studies on an *E. coli* temperature-sensitive Pth mutant found suppressor mutations that led to the overproduction of lysine tRNA, suggesting that the growth defect in *E. coli* may be due to lysine-tRNA starvation in this AT-rich organism [43]. Overexpressing other tRNAs individually, meanwhile, did not rescue the growth defect [43]. However, later work found that lysine-tRNA overexpression also led to increased protein synthesis – including Pth synthesis – raising questions on the interpretability of the lysine-tRNA suppressor and potentially pointing to a more global tRNA effect like in *Mtb* [44]. Future work investigating peptidyl tRNA pools in *pth* mutants in other bacteria would shed light on the dynamics of peptidyl tRNA drop-off and Pth’s role in other organisms. The tRNA sequencing-based methods outlined in this paper provide a scalable and efficient platform for such efforts.

Tuberculosis (TB) remains a leading infectious disease threat to global health. *Mtb* is intrinsically resistant to most existing antibiotics and current treatment requires a minimum of four months of combination therapy using potent drug regimens which have remained largely unchanged since their discovery in the mid-twentieth century. Protein synthesis is indispensable to any living cell, and in fact many of the antibiotics used to treat other bacterial infectious diseases target some aspect of protein synthesis. Nonetheless, most of these drugs are ineffective against TB, and our current anti-TB arsenal is lacking in protein synthesis inhibitors. Here, we have identified several orthogonal approaches to inhibit protein synthesis in *Mtb* by impeding an essential mediator of tRNA availability. Our work underscores Pth as a promising candidate for target-based development, given its stand-alone requirement for *Mtb* survival, as well as its ability to potentiate the activity of two classes of translation inhibitors.

Macrolides are used to treat a variety of pathogens including some non-tuberculous mycobacteria but are currently ineffective against TB. Sensitizing *Mtb* to macrolides by simultaneously targeting Pth offers a new strategy to lend efficacy to an entire class of antibiotics against this devastating disease. Many mycobacteria encode an inducible macrolide resistance mechanism [45, 46]. Exposure to macrolides triggers expression of the so-called *erm* family of genes, whose products methylate ribosomes at the site of macrolide binding. Despite possessing this innate resistance mechanism, macrolides like azithromycin are still used to treat *M. abscessus* infections, suggesting that there could still be success in including macrolides in an anti-TB regimen [35]. Since Pth depletion causes growth arrest, we hypothesize that treating Pth-depleted cells with macrolides will make it difficult – if not impossible – for cells to actively synthesize Erm methyltransferases, whose synthesis is induced by the WhiB7 intrinsic resistance transcription factor, further contributing to their sensitivity to macrolide treatment [47, 48]. Furthermore, *Mtb* clinical isolates lacking *erm* genes have been discovered in an *Mtb* sub-lineage that is endemic to Southeast Asia, and multiple other macrolide-sensitizing genes have been identified in *Mtb* and are worth exploring as additional adjuvants in a macrolide-based anti-TB strategy [47].

A recent survey of gene vulnerability to various levels of depletion using CRISPR interference (CRISPRi) found that even slightly reducing levels of tRNA synthetase expression causes major growth attenuation in *Mtb* [49]. Inhibitors against these enzymes are currently being developed, with a focus on lysine tRNA synthetases. Our work suggests that there is an opportunity to strengthen these already highly potent compounds even further by coupling them with an orthogonal regulator of tRNA availability. While existing proline tRNA synthetase inhibitors like halofuginone unfortunately do not exhibit activity against *Mtb in vitro*, our findings raise the question of whether a Pth inhibitor would synergize more strongly with proline tRNA synthetase inhibitors, given the disproportionate decrease in proline tRNA availability in a Pth knockdown.

Altogether, our work underscores the importance of Pth in tRNA turnover and provides new biological insights on anti-TB strategies that could exploit synergy between translation errors and tRNA turnover. We also demonstrate a new application of tRNA sequencing that can be used to study peptidyl tRNA and N-acetylated tRNA in other organisms, providing a high-throughput way to shed light on the dynamics of tRNA pools across all kingdoms of life.

## Supplementary Figures and Data

**Figure S1:**
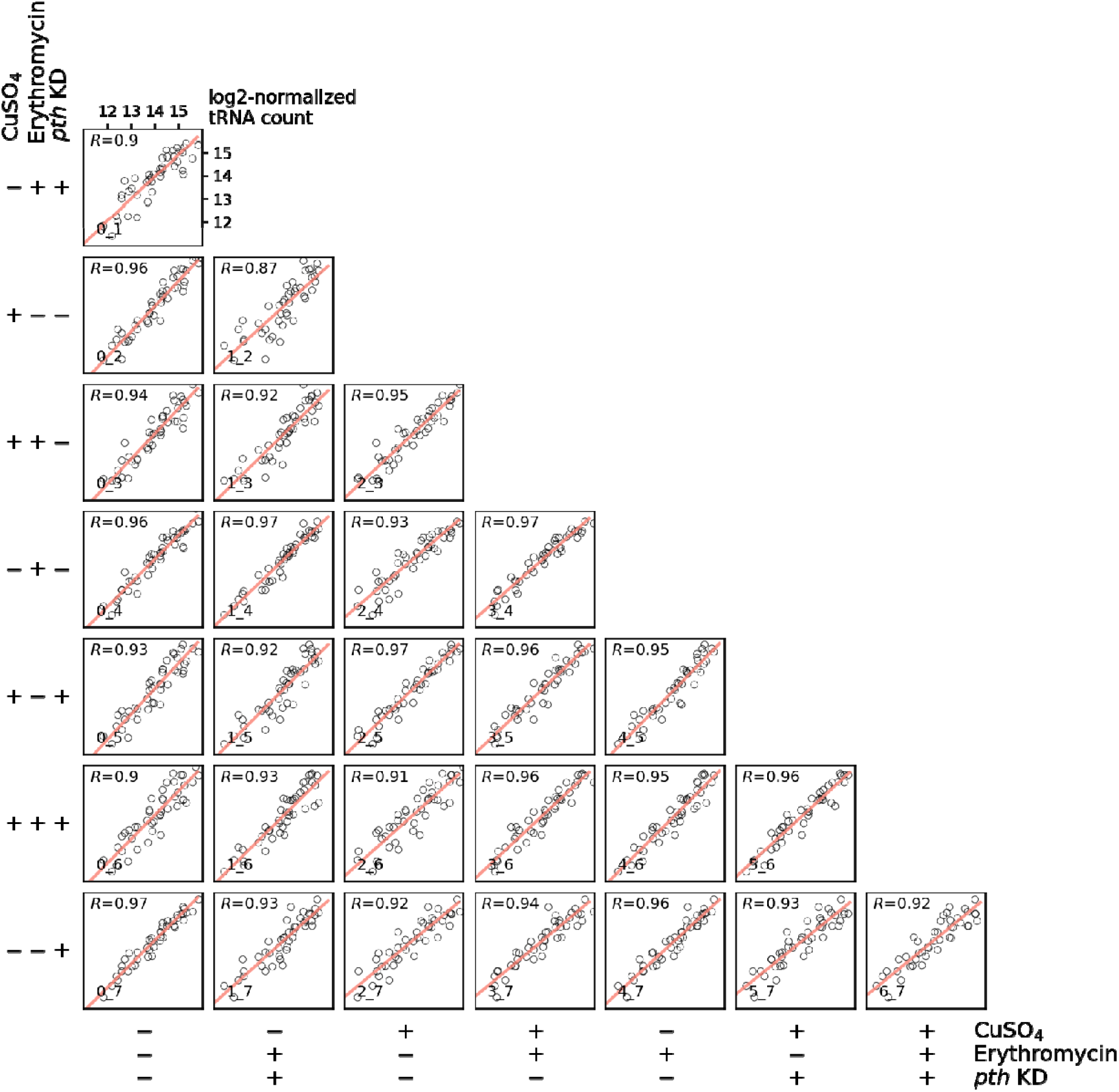
Chemical treatments do not bias tRNA counts in Cu-tRNAseq. Pairwise linear-regression and Pearson’s correlation analysis using the mean tRNA count profiles (Table S1) of different treatment groups. Pearson’s correlation coefficient is depicted on the top-left corner of each subpanel. [explain meaning of code #_# or maybe unnecessary

**Figure S2.**
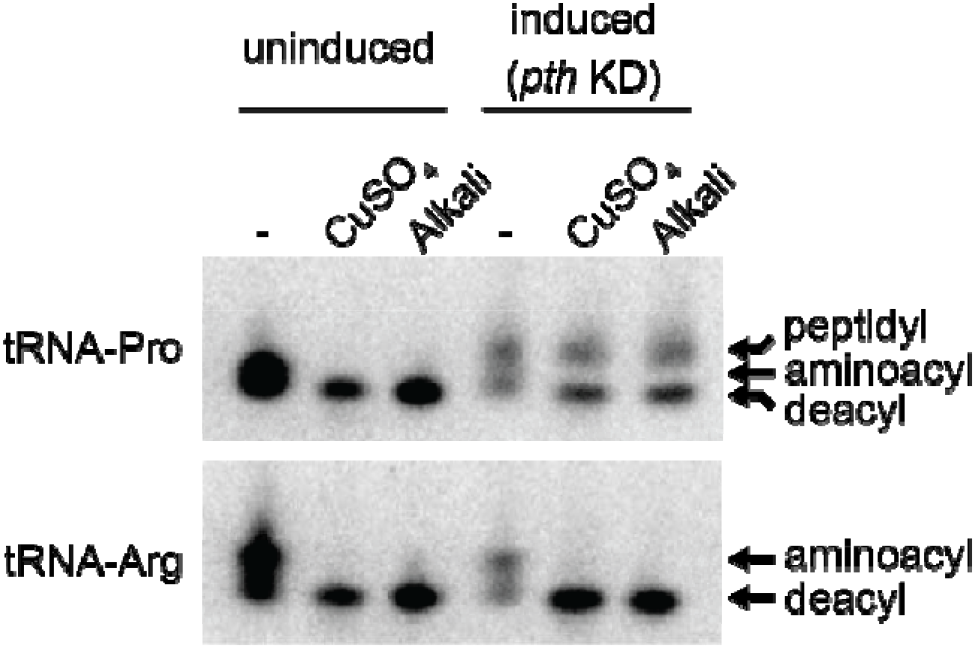
Northern blots with arginine- and proline-tRNA probes. Northern blots were performed as described in Methods. Top: RNA blotted using a probe for tRNA-Arg_ACG. Bottom: RNA blotted using a probe for tRNA-Pro_GGG. CuSO_4_ treatment was performed where indicated on total RNA samples as described in Methods. Alkali treatment, which also deacylates tRNA, was performed on total RNA where indicated (1 hour at 37°C in 100 mM Tris pH 9.0). Peptidyl-, aminoacyl-, and deacyl-tRNAs are indicated by arrows. Peptidyl prolyl-tRNA is visible in a Pth depleted condition but not in an uninduced condition. CuSO_4_ treatment deacylates aminoacyl-tRNA but not peptidyl-tRNA. KD; knockdown.

**Figure S3.**
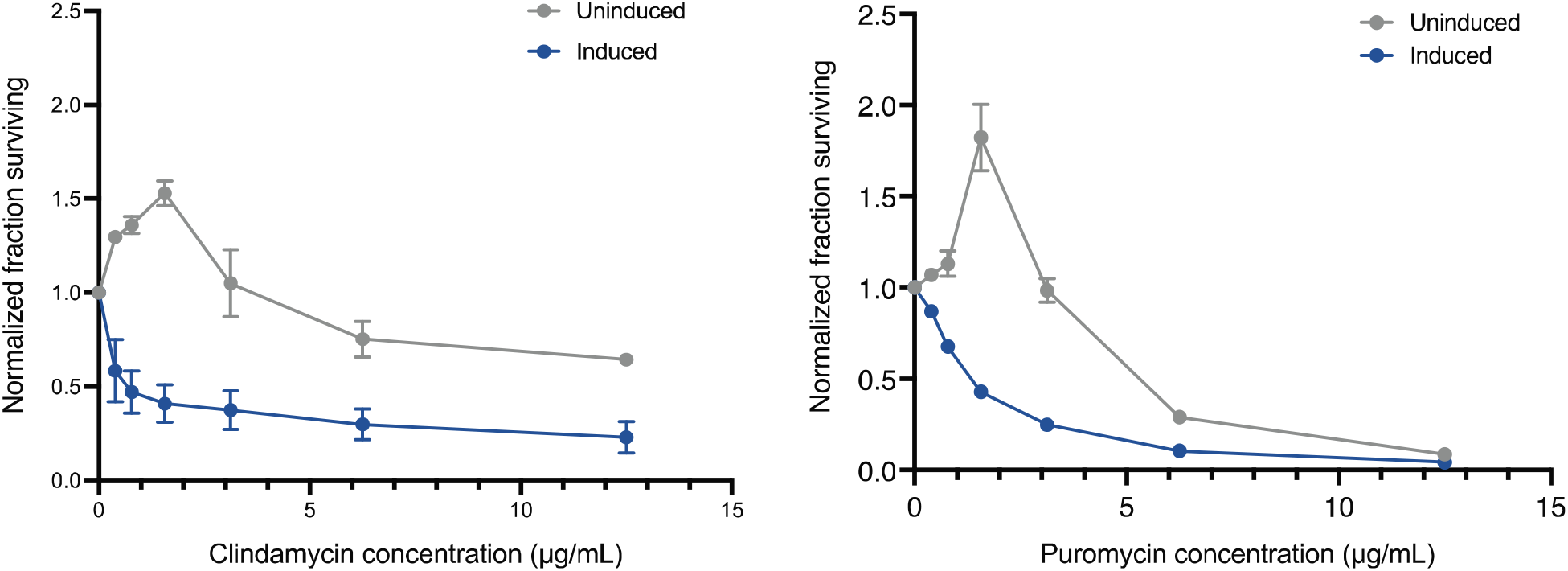
Pth depletion increases *Mtb* susceptibility to clindamycin and puromycin. *Mtb* growth in the presence clindamycin (left) or puromycin (right) is plotted across drug concentrations as determined by fluorescence in an Alamar blue assay. Pth CRISPRi strains were grown to mid-log and diluted to an OD_600_ of 0.001 and plated in serial dilutions of antibiotic in 96-well plates. The fraction of *Mtb* cells surviving is plotted by normalizing fluorescence values to control wells with no drug. Results are representative of two biological replicates.

**Figure S4:**
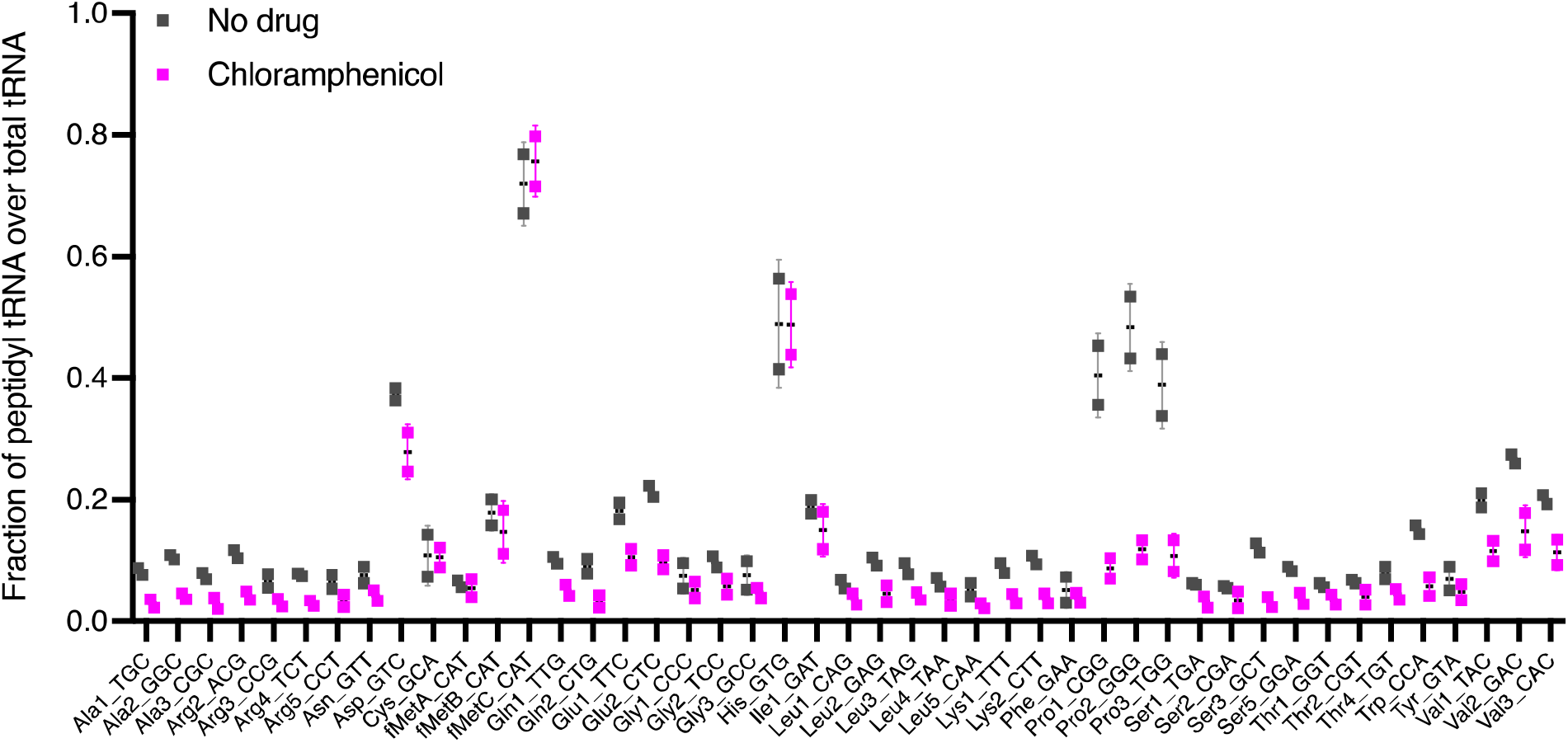
Chloramphenicol does not increase peptidyl tRNA drop-off. Cu-tRNAseq was performed on *Mtb* strains induced for Pth depletion by CRISPRi in the presence and absence of chloramphenicol (2 μg/mL). After 3 days of incubation with inducer (50 ng/mL aTC), chloramphenicol was added to cultures and incubated overnight prior to RNA extraction and sample processing. Chloramphenicol treatment (pink) led to a reduction in peptidyl tRNA pools relative to the drug-free control, suggesting that chloramphenicol-induced protein synthesis arrest affects tRNA pools differently from the other ribosome inhibitors tested in this work. Results are from two biological replicates.

**Table S2:** Minimal inhibitory concentrations of antibiotics in *Mtb* Pth proteolytic degradation strains. Minimal inhibitory concentrations are shown for a panel of antibiotics against Pth depletion strains by proteolytic degradation. Pth-tagged control: *pth-flag-DAS* strain used as a parental backbone for generating knockdown strains (by introducing inducible SspB expression plasmids). This strain serves as a control for potential effects of the FLAG or DAS tags on Pth activity even in the absence of induced degradation mediated by SspB.

|  | Pth-tagged control | pTetON-6 (-) | pTetON-6 (+) | pTetON-10 (-) | pTetON-10 (+) |
| --- | --- | --- | --- | --- | --- |
| **Isoniazid** | 0.06 | 0.06 | 0.06 | 0.06 | 0.06 |
| **Ethambutol** | 0.59 | 0.59 | 0.59 | 0.59 | 0.59 |
| **Kanamycin** | 1.56 | 2.34 | 2.34 | 2.34 | 2.34 |
| **Chloramphenicol** | 1.2 | 0.59 | 0.59 | 0.59 | 0.59 |
| **Fusidic acid** | 0.30 | 0.30 | 0.30 | 0.30 | 0.30 |
| **Linezolid** | 0.15 | 0.15 | 0.15 | 0.15 | 0.15 |
| **Capreomycin** | 1.56 | 0.78 | 0.78 | 1.56 | 0.78 |
| **Erythromycin** | 6.25 | 0.4 | 0.05 | 0.4 | 0.012 |
| **Clarithromycin** | 5 | 0.08 | 0.02 | 0.04 | < 0.01 |

**Table S3.**
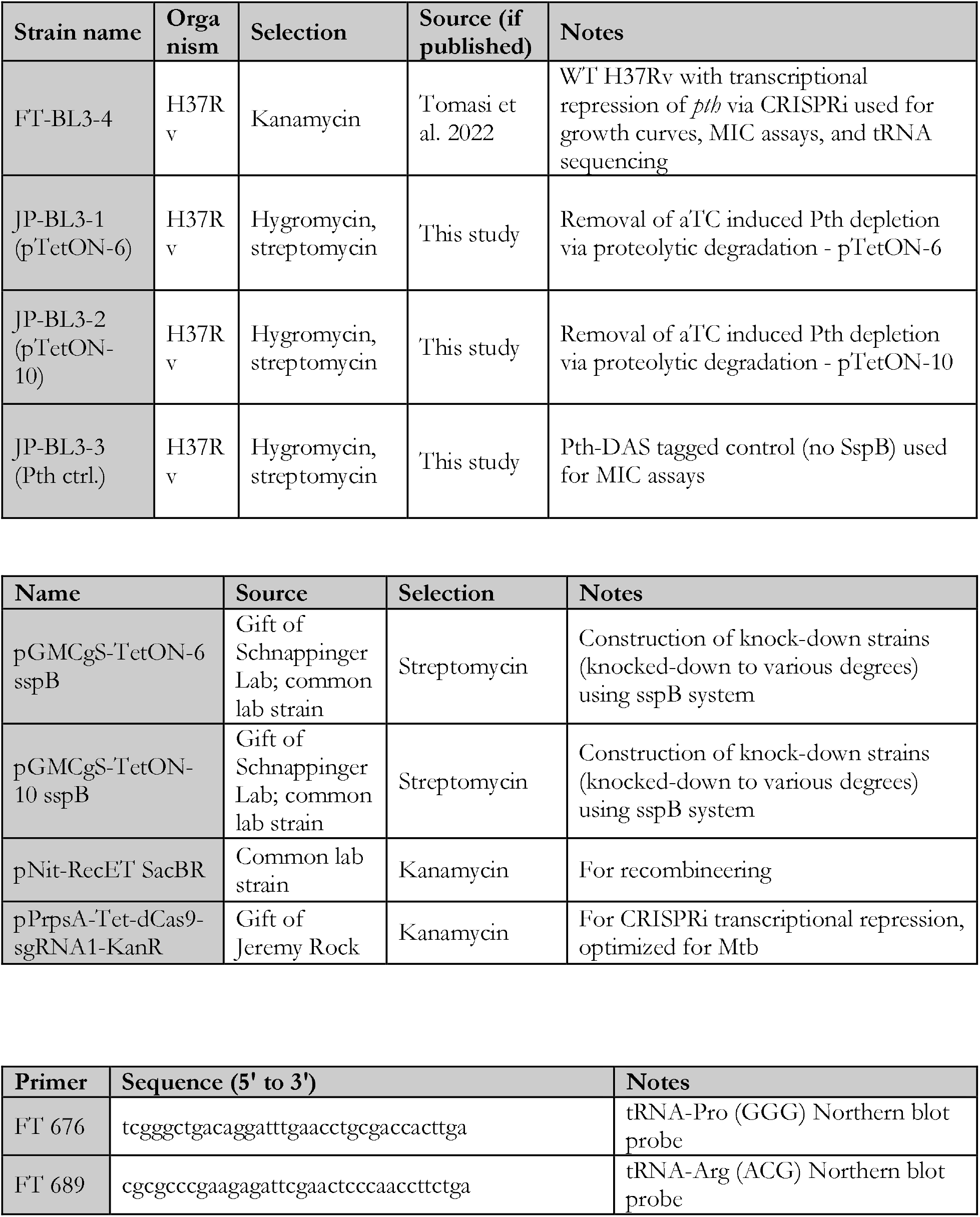
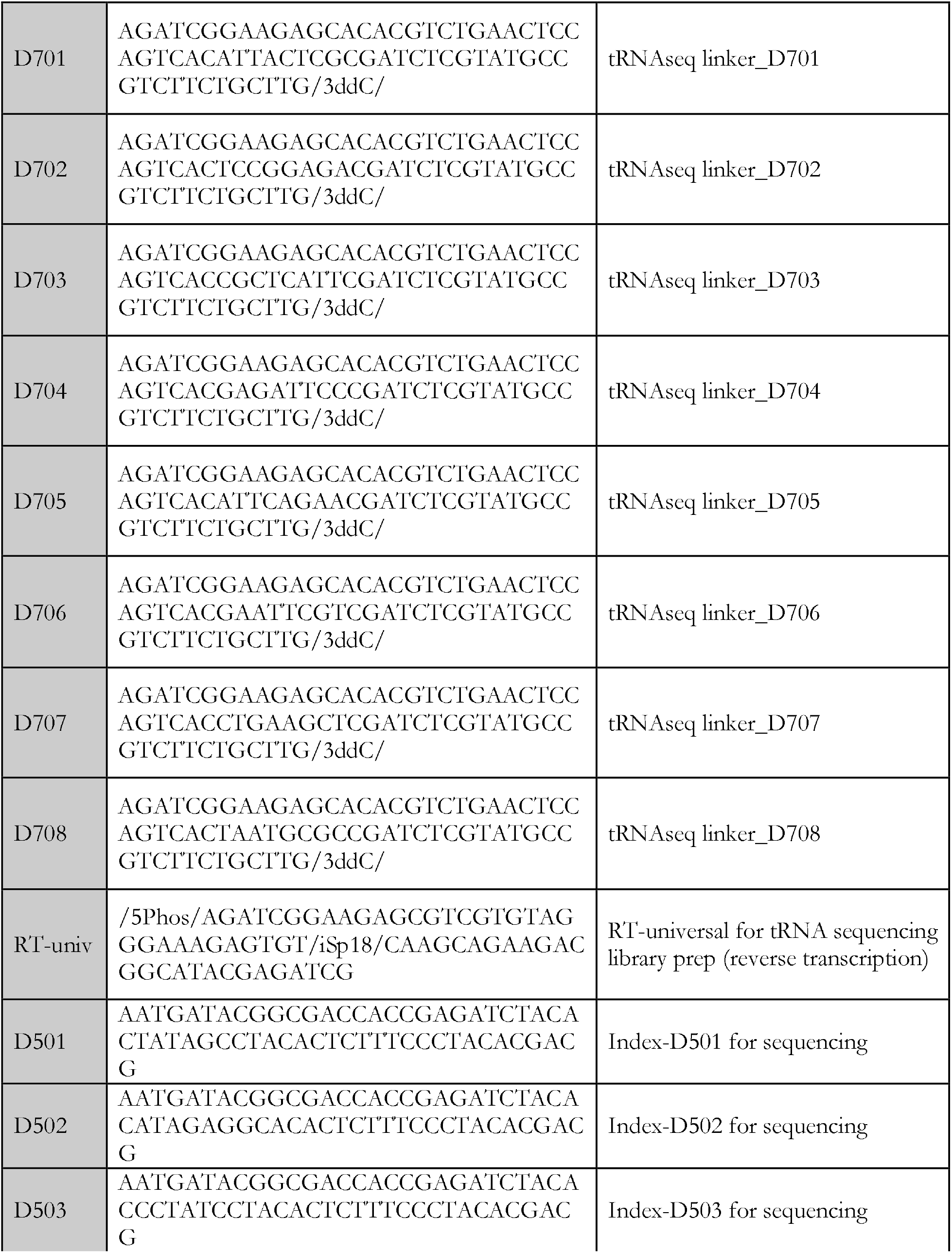
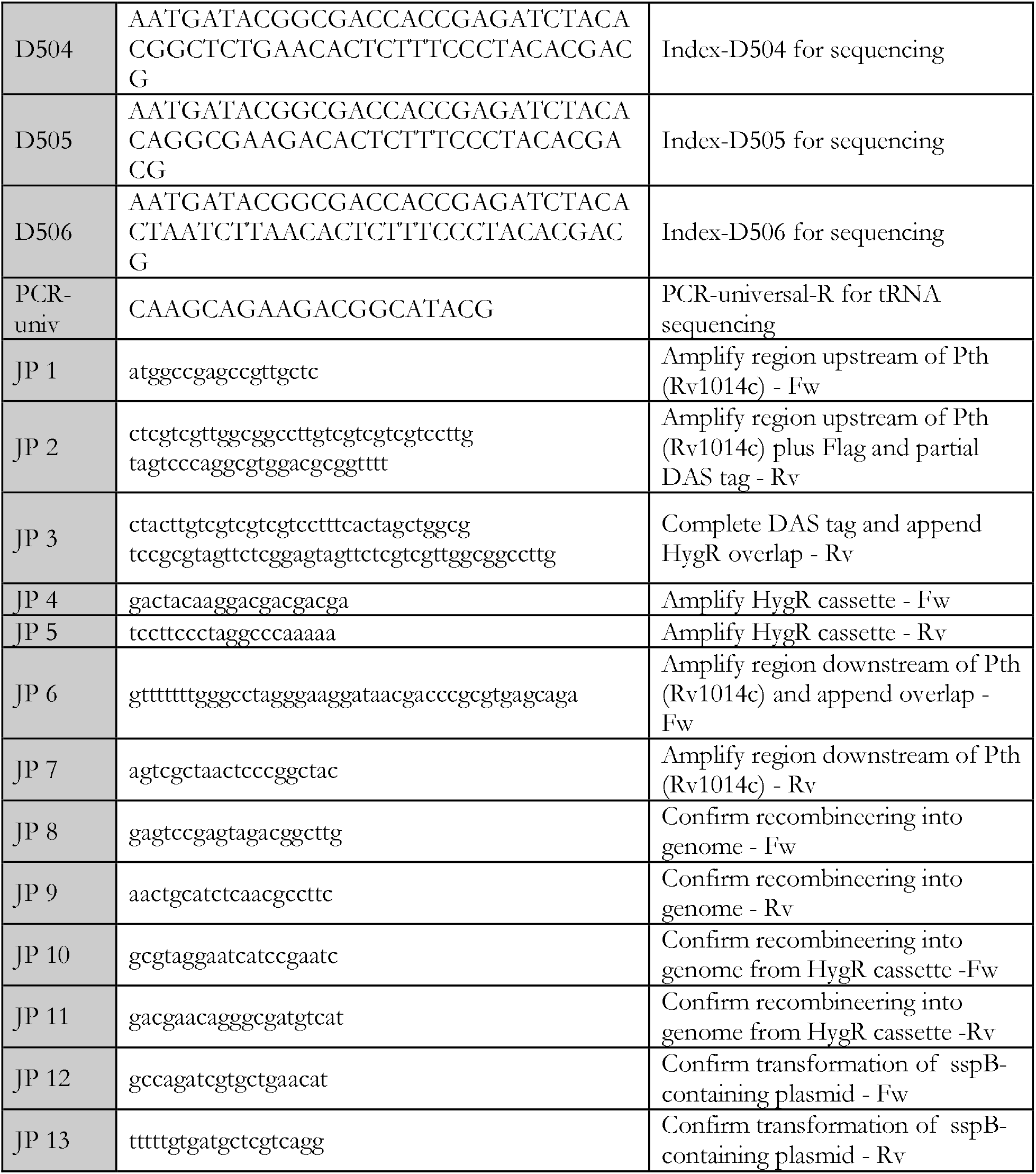
Strains and primers used in this work. All strains and primers used for the work in Chapter Three are listed below. Top; strains. Middle; plasmids. Bottom; primers

## Materials and Methods

### Bacterial strains and growth conditions

*Mtb* and *Msmeg* strains were grown from frozen stocks into Middlebrook 7H9 medium supplemented with 0.2% glycerol, 0.05% Tween-80, and ADC with or without oleic acid, respectively (5 g/L bovine serum albumin, 2 g/L dextrose, 3 μg/ml catalase). Cultures were incubated at 37 °C. Antibiotics or inducing agents were used when needed at the following concentrations in both *Mtb* and *Msmeg*: kanamycin (25 μg/ml), anhydrous tetracycline (aTC; 100 ng/mL), hygromycin (50 μg/ml), and nourseothricin (20 μg/ml). Transformed *Mtb* and *Msmeg* strains were plated onto 7H10 agar plates with the appropriate antibiotic(s). Strains were grown to mid log-phase for all experiments unless otherwise specified (OD_600_ 0.4-0.6). *E. coli* strains for cloning or protein purification were grown in LB broth or on LB agar with appropriate antibiotics at the following concentrations: kanamycin (50 μg/ml), zeocin (50 μg/ml), and nourseothricin (40 μg/ml). Induction time for *pth* depletion in *Mtb* was 3 days.

### Bacterial strain construction

Table S3 depicts the strains, plasmids, primers, and recombinant DNA used for this study. Plasmids were built by restriction digest of a parental vector and inserts were prepared either by restriction enzyme cloning or Gibson assembly using 40 bp overhangs, as specified in Table S3. Plasmids were isolated from *E. coli* and confirmed via Sanger sequencing carried out by Genewiz, LLC (Massachusetts, USA).

#### Pth-knockdown constructs

Transcriptional knockdown of *pth* was accomplished using mycobacterial CRISPRi-interference (CRISPRi) as described with the same strain used in Chapter Two [22]. Proteolytic degradation strains were built using the plasmids and primers listed in Table S3. Briefly,

### Northern blotting

Acid-PAGE Northern blotting of tRNA was performed as previously described [50]. Briefly, 0.5 μg total RNA was electrophoresed on a 6.5% urea gel (7 M urea, 100 mM NaOAc, pH 5.0), transferred to a nitrocellulose membrane by semi-dry blotting, and UV-crosslinked twice (1200 μJ). Membranes were incubated with ULTRAhyb-oligo (ThermoFisher Scientific) at 42 °C for 30 min followed by hybridization overnight at 42°C with 4 pmol probes (Table S3) that were radiolabeled using [γ^-32^P] ATP (PerkinElmer) and T4 Polynucleotide kinase (New England Biolabs). Membranes were washed twice with 2xSSC (3M NaCl, 300 mM sodium citrate 2H_2_O) + 0.1% SDS and bound probe was detected using a FLA-5000 PhosphoImager (Fuji).

### tRNA sequencing

#### Extraction of total RNA

Strains were grown to mid-log phase with the appropriate antibiotics and inducing agents described above. RNA was collected at the same OD_600_ for each strain (between 0.4–0.6). Cells were left on ice for 20 minutes, then pelleted by centrifuging at 4,000 rpm for 10 minutes at 4C. Pellets were resuspended in 0.5-1 mL of TriZol (Life Technologies) and lysed using a BeadBug microtube homogenizer (Millipore Sigma). 200 μL of chloroform was added to each tube, after which samples obtained from *Mtb* strains were removed from biosafety level 3 precautions. Samples were centrifuged at 15,000 rpm for 15 minutes at 4°C and the aqueous layer was collected into a fresh tube. To the original tube, 250 μL of sodium acetate buffer (300 mM sodium acetate pH 5.2 and 10 mM EDTA pH 8.0) was added, and samples were vortexed at 4C for 5 minutes then centrifuged at 15,000 g for 15 minutes at 4°C. The aqueous layer was added to the fresh sample-containing tubes. 400 μL chloroform was added, and tubes were briefly vortexed and then centrifuged at 15,000 rpm for 1 minute at 4°C. The aqueous phase was collected into a fresh tube and RNA recovered by ethanol precipitation. RNA pellets were resuspended in 10mM sodium acetate pH 5.2 and stored at −80°C until processed for sequencing.

#### Copper sulfate treatment

Copper sulfate treatment of total RNA samples was performed as described [23]. Briefly, 20-30 μg of total RNA was added to fresh tubes to a final volume of 54 μL in storage buffer (8 M urea, 10 mM sodium acetate pH 5.2, 1 mM EDTA). 6 μL of 100 mM copper sulfate (CuSO_4_) were added for a final concentration of 10 mM CuSO_4_ and reactions were incubated at 37°C for 1 hour. 1mM EDTA was then added to each tube, along with 3 μL of glycogen (2 mg/mL). Samples were recovered by ethanol precipitation and stored at −80°C between treatments.

#### Sodium periodate treatment

Sodium periodate (NaIO_4_) treatment of total RNA samples was performed as described [27]. Briefly, 1 μg of total RNA was combined with 100 mM sodium acetate pH 5.2 and 50 mM NaIO_4_ in a final volume of 100 μL. Reactions were incubated at room temperature in the dark for 30 minutes, and subsequently quenched with 100 mM glucose for 5 minutes. Unquenched periodate was removed using Micro Bio-Spin P-6 columns (Bio-Rad Laboratories) according to manufacturer instructions. RNA was recovered by ethanol precipitation.

#### Beta-elimination treatment

Beta elimination of periodate-oxidized RNA samples was performed as described [27]. Briefly, total RNA collected after periodate oxidation was combined with 60 mM sodium borate pH 9.5 (Boston BioProducts) in a final volume of 100 μL and incubated for 90 minutes at 45°C. Samples were purified with Micro Bio-Spin P-6 columns (Bio-Rad Laboratories) according to manufacturer instructions. RNA was recovered by ethanol precipitation.

##### Library preparation

###### Isolation of tRNA fraction

1-2 μg of total RNA was run on a 10% TBE-UREA gel (ThermoFisher Scientific) at 250 V for 1 hour. Gels were stained with SYBR Gold (ThermoFisher Scientific), and tRNA was excised. Excised gels containing tRNA fractions were mashed in RNAse-free tubes, and 300 μL elution buffer (300 mM NaOAc pH 5.5, 1 mM EDTA pH 8.0, 0.10% SDS) was added to each tube. Samples were shaken on a thermoshaker (VWR) for 1-4 hours at 37°C and supernatant was collected using an Ultrafree filter column (Millipore Sigma). tRNA was recovered by isopropanol precipitation.

###### tRNA dephosphorylation

tRNA was dephosphorylated using QuickCIP (New England BioLabs) according to manufacturer instructions, and tRNA was collected by phenol-chloroform extraction followed by isopropanol precipitation.

###### Adapter ligation

0.5 μL RNase inhibitor was added to 3.5 μL dephosphorylated tRNA (200-250 ng tRNA) and samples were boiled at 80°C for 2 minutes. Boiled tRNA was mixed with 12 μL PEG buffer mix (10 μL 50% PEG8000, 2 μL 10 × buffer B0216S; New England Biolabs). 3 μL of 5’ adenylated linkers (Table S3) were added (33 pmol/μL) along with 1 μL T4 RNA ligase 2 truncated (New England BioLabs) and incubated at 25°C for 2.5 hours. Samples were recovered by isopropanol precipitation and run on a 10% TBE-Urea PAGE gel for 40 minutes at 250 V. Ligated products were recovered by gel excision as described above.

###### Reverse transcription

Identical quantities of samples with different adapter sequences were pooled for reverse transcription for a total of 200-250 ng tRNA. Reverse transcription was performed by combining 2.1 μL dephosphorylated tRNA with 100 mM Tris-HCl pH 7.5, 0.5 mM EDTA, 1.25 μM RT primer (Table S3), 450 mM NaCl, 5 mM MgCl_2_, 5 nM DTT, 500 nM TGIRT (InGex), and 15% PEG8000 in a final volume of 9 μL. Samples were incubated at 25°C for 30 minutes, after which 1 μL 10 mM dNTPs (New England BioLabs) were added and reactions incubated at 60°C for 1 hour. 1.15 μL NaOH was added, and samples were boiled for 15 minutes and run on a 10% TBE Urea PAGE gel at 250 V for 1 hour. Reverse transcription products were excised from and cDNA was recovered by isopropanol precipitation. Linear single stranded cDNA was circularized using CircLigase II (Lucigen) in accordance with manufacturer instructions.

###### PCR of tRNA libraries

PCR reactions were set up using HF Phusion according to the manufacturer’s instructions using a universal reverse primer (Table S3) and a different index primer for each pool of samples. PCR reactions were aliquoted into 4 tubes and collected after 6, 8, 10, and 12 cycles. Samples were run on a Native TBE PAGE gel (ThermoFisher Scientific) at 180 V for 50 minutes, and amplified products were cut from the same cycle for each sequencing run. Samples were recovered by gel excision and isopropanol precipitation.

###### Sequencing

Sequencing was performed on a MiSeq instrument (Illumina) using 150 bp single end reads with a version 3, 150 cycle kit.

###### Analysis

3’ linker sequences and two nucleotides at the 5’ end were trimmed using cutadapt and fastx-trimmer. Bowtie v1.2.2 was used with default settings to map reads to reference Mtb or Msmeg tRNA sequences (Datasets S1 and S2) retrieved from Mycobrowser [51]. Mpileup files were generated using samtools (samtools mpileup -I -A --ff 4 -x -B -q 0 -d 10000000). 3’ end termini of mapped reads were piled up using the bedtools genomecov command (-d -3 -ibam). To analyze 3’ ends, a cutoff of 500 read counts for each tRNA species was set. The number of 3’ termini at any position was divided by the total number of mapped termini to compute 3’ termination frequency. The ratio of peptidyl tRNA for each tRNA species is the 3’ termination frequency at the terminal A residue of the 3’ CCA tail of each tRNA sequence compared to the sum of the frequencies at the terminal A and at the previous C nucleotide.

### Drug susceptibility assays

Drug susceptibility was measured using a minimum inhibitory concentration (MIC) assay as described [39]. Briefly, strains were diluted to an OD_600_ of 0.001 and tested in technical duplicate in serial dilutions of the following antibiotics: erythromycin (GoldBio), clarithromycin (GoldBio), puromycin (GoldBio), clindamycin (GoldBio), chloramphenicol (Sigma Aldrich), isoniazid (Sigma Aldrich), ethambutol (Sigma Aldrich), kanamycin (IBI Scientific), fusidic acid (Sigma Aldrich), linezolid (Sigma Aldrich), and capreomycin (Sigma Aldrich). Lysine tRNA inhibitors were synthesized at the University of Dundee (in preparation) and dissolved in DMSO. When needed (ex. for *pth* knockdown and empty guide RNA controls), strains were pre-induced with aTC as described above. Each MIC assay was conducted in biological triplicates, with each biological replicate conducted in technical duplicate. In plates containing CRISPRi induced cells, aTC was added to each well at a final concentration of 100 ng/mL. Here, a technical replicate is considered a single row of drug and bacterial incubation, using bacteria from the same culture. Biological replicates are considered plates set up in the same way using bacteria from a different culture. 96-well plates were agitated at 37°C for 6 days, at which point 0.0002% resazurin was added and plates were agitated at 37°C for an additional 48 hours. Plates were read on a BioTek plate reader at both 24 hours and 48 hours. Resazurin was measured by OD_570_, and OD_600_ was also measured for each well. Fluorescence was normalized to OD_600_ and to the positive control for each strain (no drug; in CRISPRi induced wells, the positive control still contained aTC), and the fraction of bacteria surviving relative to the no-drug control well was plotted against drug concentration for each well.

## Notes

### Competing Interest Statement

The authors have declared no competing interest.

